# Nutrient control enables metabolic reconstruction of *L. rhamnosus* GG and analysis of secretions

**DOI:** 10.64898/2026.06.02.729517

**Authors:** Gracelyn R. Richmond, Emanuel Cunha, Libusha Kelly, Óscar Dias, Roger L. Chang

## Abstract

*Lacticaseibacillus rhamnosus* GG (LGG) is an important gut commensal bacterial strain that has been extensively studied in both industrial and health settings. Despite its long history of study, a high-quality genome-scale metabolic network model (GEM) for LGG has yet to be reconstructed. Only automatically-generated draft models have been published, which have notoriously limited functional accuracy. Furthermore, comprehensive nutrient requirements have not been established for well-controlled *in vitro* study. Here we present the first curated GEM for LGG using a new approach for reconstruction and validation that leverages multiple automatically-generated draft models, applied study literature, and high-throughput defined media experiments. In addition, our results include a series of chemically defined media, extensive single-component nutrient dropout growth data, insights from *in silico* and *in vitro* experiments into major secretion products lactate and indole-3-carboxaldehyde, a minimal medium and *in silico* characterization of LGG’s nutrient requirements. Our approach for developing interdisciplinary research tools for LGG metabolism comprises a new framework that could be applied to many understudied microorganisms, particularly useful in studying bacteria within the human microbiome.

## Introduction

*Lacticaseibacillus rhamnosus* GG (LGG) is an important gut commensal bacterial strain that has been studied in food microbiology and probiotic development since the 1980s^1^. LGG has been studied in the context of a variety of host health conditions, including alleviation of inflammatory bowel disease (IBD) symptoms and gastrointestinal (GI) side effects of medical treatments^2–4^ and for improving mental health and neurodegenerative conditions^5–7^. LGG is known to secrete many small molecule metabolites with beneficial host effects and participating in interactions with other members of the gut microbiota. For example, LGG is widely known to secrete lactate as a product of heterolactic fermentation, its primary mode of anaerobic energy generation. Secreted lactate becomes available as a metabolic substrate for other members of the microbiome, transforming into host-beneficial bacterial metabolites through metabolic cross-feeding^8–11^. LGG itself can also produce host-beneficial metabolites, such as indole-3-carboxaldehyde (I3A)^12^. I3A and other tryptophan degradation products have been shown to alleviate host tight junction stress^4,13–15^. I3A and LGG gavage both improve overall survival during gut stress and I3A is enriched in healthy guts experiencing stress^16^. However, I3A presents a special challenge to study and leverage within the microbiome due to incomplete knowledge of its synthetic pathway^15,17–20^.

Despite the importance of LGG in microbiome-based interventions, critical tools are lacking for mechanistic and systematic *in vitro* and *in silico* study of its metabolism. In the age of metagenomics, many bacteria identified as important biomarkers or with therapeutic or industrial potential have primarily been studied in an applied manner^18^; such is the case for LGG studies, which mostly explore application in amelioration of disease states^21–24^. However, findings from such studies are not often expanded upon mechanistically, leaving much of the state of knowledge about important bacteria correlative and scattered across systems and applications. These translational applications will remain limited without a fundamental understanding of LGG metabolism. Development of tools, like *in silico* reconstruction of genome-scale metabolic network models (GEMs), would provide such a fundamental foundation.

Traditionally, high-quality GEM reconstruction is a largely manual four step process that involves: 1) building a draft model of candidate reactions and genes that are supported by genome functional annotation, 2) refinement of the model to ensure basic functionality and biological completeness, 3) transforming the model into a computable format, and 4) evaluating the model for missing content and testing known capabilities of the organism through flux simulations^25^. This approach requires extensive biochemical knowledge, literature, and experimental data for the subject organism. These requirements make curation of a GEM difficult for bacteria like LGG, which does not have a wealth of mechanistic studies available.

More recently, automated methods based on genome functional annotation have been developed to perform much of steps 1, 2, and some extent of refinement in step 3^26–29^. Automated approaches were recently applied to generate draft models for thousands of taxa from the human microbiome^26,30^, and one study has used an LGG GEM derived in this manner^31^. However, such automatically-generated reconstructions are limited by the completeness, accuracy, and cross-referencing of source databases, algorithm-specific functionality corrections, and lack of refinement through literature curation, experimental validation, and model testing^32–34^. A previous meta-analysis that compared 7 different automated approaches outlined their strengths and limitations and suggested that field standards must be developed to ensure biological accuracy of reconstructions, rather than just functional outputs^34^. Automatically-generated draft reconstructions are not expected to be fully functional or as biologically accurate as traditionally reconstructed, manually-curated GEMs. However, the advancing resource landscape offers opportunities to further improve upon automated approaches and develop protocols for reconstruction that leverage experimental information.

Previous high-quality, manually-curated GEMs relied heavily on high-throughput genetic screens and mutant libraries to enable step 4. For mechanistically understudied bacteria such as LGG, genetic tools and even fully-controlled culture methods have been lacking, challenging high-quality GEM reconstruction. Previous culture protocols for LGG do not even enable full nutritional control, such as a chemically-defined minimal medium, and the limited tools developed for other organisms and applied to LGG are not widely used^35^. Not only does nutritional control using defined media improve the reproducibility and rigor of mechanistic biochemical and microbiological studies, but it also enables high-throughput *in vitro* experiments to provide rich metabolic phenotype datasets, analogous to a mutant library, for GEM evaluation and refinement. In addition, the knowledge gained in establishing nutritional control and a high-quality GEM can also provide a systems-level mechanistic definition of an organism’s metabolic niche, defined by its environmental nutrient requirements and the range of its metabolite secretion products.

By bridging mechanistic frameworks for GEM reconstruction and achieving nutritional control *in vitro* and *in silico* in this study, we have broadly expanded the understanding of LGG metabolism. Here, we introduce a series of chemically-defined media for LGG with varying nutritional complexity, a high-throughput single-component dropout experimental growth phenotype screen, a manually-curated GEM called *i*GR384, a minimal medium paired with *in silico* characterization of nutritional requirements and their contributions to biomass component production, and *in vitro* and *in silico* characterization of the key secreted metabolites lactate and I3A. Together, these results provide new practical and theoretical tools for the study of LGG metabolism and exemplify a framework for the reconstruction of GEMs for bacteria with similarly limited mechanistic depth of published studies.

## Results

### Consensus approach to generate high-confidence reconstruction of LGG metabolism

Our reconstruction protocol for understudied microbes builds upon the traditional protocol and introduces some new strategies (Figure 1A). Our method includes merging multiple independently-derived, automatically-generated draft reconstructions (Step 1.5) by namespace mapping metabolites, reactions, and genes to unify shared components and addition of unique components from each source reconstruction. Across Steps 2, 3, and 4, we iteratively prune and add reactions to the reconstruction considering *in vitro* growth phenotype data. This data comes from extensive single-component dropout experiments from chemically defined media. Traditional reconstruction validation and curation often rely on high-scale single-gene deletion data, while our approach uses a similar logic for environmental nutrients rather than genes. Our approach integrates *in vitro* phenotypic experiments and *in silico* growth simulations to identify model gaps in functionality to be filled. Completion of this reconstruction process for LGG yielded *i*GR384 (Supplemental Files 1-2), which accounts for activity of 384 gene products. Information regarding curation decisions is reported in Supplemental File 2.

**Figure 1.**
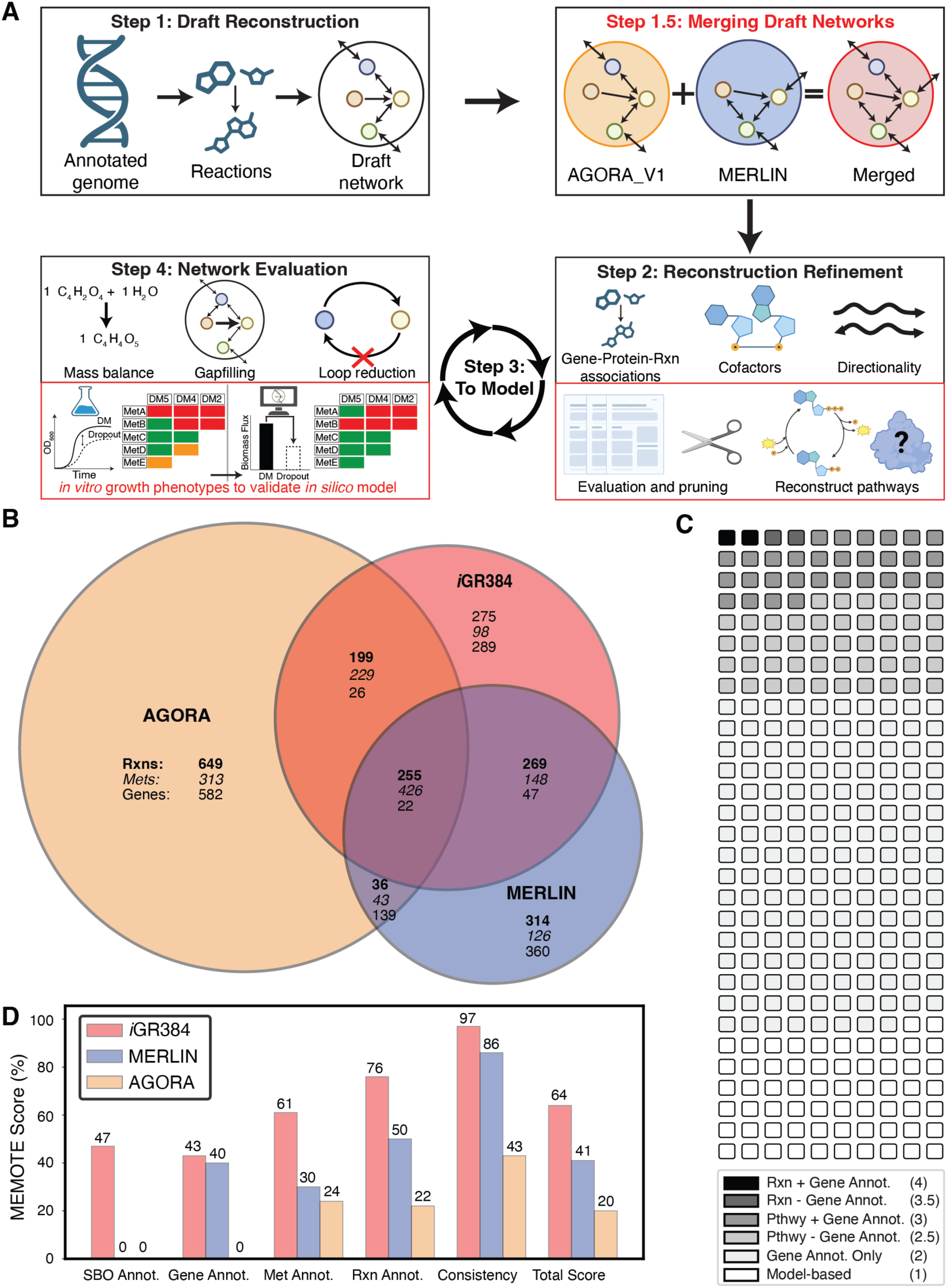
Reconstruction of *i*GR384 and summary of models. *A. Reconstruction overview.* Red text indicates approaches introduced in this study. *B. LGG model content comparison*. Numbers of shared and unique reactions (bold), metabolites (italics), and genes across models are given. *C. iGR384 reaction evidence levels.* Each patch represents 5 reactions with a given evidence level (see Supplement Evidence Table). *D. MEMOTE evaluations for LGG models.* MEMOTE score as calculated using MEMOTE version 0.17.0 differential report with default parameters.

While about 72% of *i*GR384’s 997 reactions derive from the AGORA and MERLIN (Supplemental File 3) reconstructions, about 28% represent new content based on literature and curation based on observations from *in vitro* experiments in this study (Figure 1B). The reconstruction process involved significant pruning of erroneous or poorly supported content from the AGORA and MERLIN reconstructions. Major error types include improper assignment of gene annotation to reactions without evidence, literature evidence suggesting pathways not present in LGG or close relatives, reaction and metabolite identifiers that were not included in standard databases (KEGG^36^, BioCyc^37^, BiGG^38^, BRENDA^39^, and MetaNetX^40^), and non-gene-associated blocked reactions that existed only to enable flux through previously pruned reactions. For the AGORA reconstruction, this process resulted in 57%, 29%, and 76% of reactions, metabolites, and genes being removed, respectively (Supplement File 4). For the MERLIN reconstruction, 40%, 22%, and 88% of reactions, metabolites, and genes were removed, respectively (Supplemental File 4). Additionally, 40 applied studies were identified and used in the reconstruction, expansion, and refinement of *i*GR384.

Our reconstruction process involved a modified reaction evidence scoring system adapted from the standard reconstruction protocol^25^ (Supplemental Table 1) to account for the dearth of mechanistic studies on LGG metabolism. Only 27% of reactions in *i*GR384 have stronger evidence than simple gene functional annotation (Figure 1C); only 13 reactions had direct, reaction-specific evidence from literature (Supplemental File 2), highlighting the need for more mechanistic studies of LGG metabolism to gain more complete knowledge and subsequently improve the reconstruction.

### *i*GR384 has higher annotation and topological quality than automatically-generated draft reconstructions

We comparatively evaluated the topological and annotation quality of *i*GR384 and the draft reconstructions to ensure completeness, usability, and to highlight limitations using the standardized MEMOTE tool^41^. *i*GR384 has a total MEMOTE score of 64% while the AGORA and MERLIN reconstructions scored 20% and 41% respectively, demonstrating substantial improvement with *i*GR384 (Figure 1D, Supplemental File 2). *i*GR384 has the highest score in all MEMOTE categories. Additionally, MEMOTE topology tests reveal that all reaction directionalities in *i*GR384 are accurate and with no missing cofactors, and all transporter reactions are defined correctly. All three reconstructions have no thermodynamically infeasible loops, enabling easily interpretable flux states using the most basic simulation tools. Loops can lead to artificial flux inflation in flux balance analysis (FBA) simulations, masking thermodynamically accurate quantitative flux values^42^. Generally, *i*GR384 is better annotated and has strong topological consistency compared to the draft models, indicating its greater utility for future applications.

### Nutrients supporting LGG growth characterized by iterative single-component dropouts in defined media

Precise nutrient control is required to enable detailed characterization of the range of LGG metabolic capabilities and as part of our network reconstruction and validation strategy. We developed a rich defined medium (DM) with 52 components (DM52) (Supplemental File 5) by combining formulations of three different chemically defined media for *Lactobacilli*^43–45^. Iterative reduction of DM52 via single-component dropout experiments yielded a series of chemically defined media (Figure 2A). These media formulations (DM52, DM24, DM16, and DM13) vary in complexity and functional redundancy. LGG grows in all media formulations with differing growth characteristics (Figure 2B). Dropouts with decreased growth rates indicate components required for maximal growth. More minimal media in the series generally amplified the negative effects of individual dropouts that always have a deleterious effect (e.g. cysteine) and those with no initial effect in DM52 (e.g. aspartate). Each defined medium includes compounds that satisfy basic growth requirements including energy sources, carbon sources, nitrogen sources, amino acids, nucleotides, mineral salts, and vitamins/cofactors (Figure 2A). As the media become more minimal, redundancy in each growth requirement decreases. Contrary to expectation, the richest medium DM52 does not support the highest growth rate of the series (Figure 2B), due to the presence of 17 compounds with statistically significant increases in growth rate when removed, suggesting an inhibitory effect.

**Figure 2.**
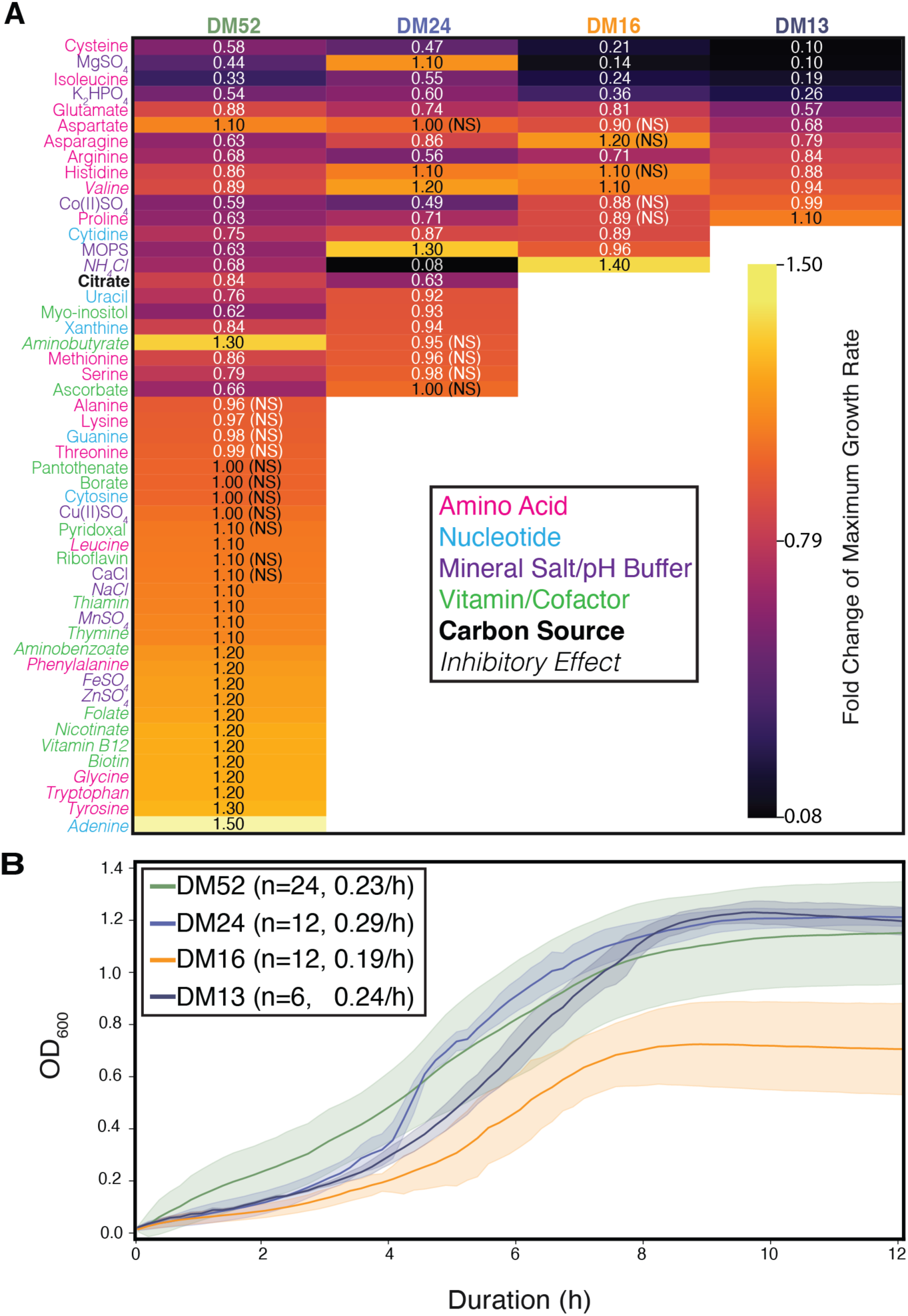
Development of defined media (DM) series for LGG culture by single-component dropouts and associated growth data. *A. Effects of single-component dropouts in DM formulations*. Fold changes of average maximum specific growth rate across six technical replicates per experiment compared to respective complete DM formulations are shown. Components are sorted by ascending fold change first in DM52, then DM24, DM16, and DM13 in turn. Empty boxes in more minimal subsequent formulations represent components removed due to inhibitory or no effect on growth. Component labels are color-coded by nutrient requirement categories. All fold changes were statistically significant (unpaired t-test p-value < 0.05) unless otherwise noted as NS (not significant). *B. LGG growth curves in DM series.* Average OD600 measurements are shown with standard deviations with a 10-minute measurement interval for all DM formulations. The number of technical replicates and average maximum specific growth rate for each DM are listed in the inset key.

These negative growth effects likely indicate activation of feedback regulatory mechanisms. Indeed, many of these regulatory mechanisms have been elucidated in other bacterial species, including in other lactic acid bacteria (LABs). High concentrations of phenylalanine, glycine, tryptophan, and tyrosine, along with neutral to slightly alkaline conditions, as in the DM series, results in L-amino acid conversion to D-amino acids. These enantiomers have been shown in *Staphylococcus epidermidis, Lactobacillus cellobiosus, Lactobacillus plantarum,* and *Bacillus subtilis* to be improperly incorporated into peptidoglycans and inhibit growth^46–48^. Additionally in *E. coli* K-12, excessive adenine has been shown to disrupt the balance of purine nucleotides and subsequently inhibit growth^49^. In *Lactobacillus sake* and *Lactobacillus curvatus,* it has been shown that sodium chloride can induce osmotic stress and inhibit growth^50^. Zinc has also been observed to inhibit growth of *Lactobacillus acidophilus* and *L. plantarum*^51^. The inhibitory effect of valine has been well-characterized in *E. coli* K-12, where it plays a role in both regulation of the *ilvB* operon and can form inhibitory complexes with isoleucyl-tRNA synthetase and reduce isoleucine levels^52–55^.

The resulting series of defined media exhibit average maximum growth rates in the range of 0.15/h to 0.29/h, with the greatest rate achieved in DM24 (Figure 2B). Additionally, growth curve features are generally conserved between these defined media and the chemically-undefined rich MRS broth previously published, which boasts a max growth rate of ∼0.27/h^56^.

The increased growth rate achieved moving from DM52 to DM24 is likely explained by the cumulative elimination of 17 seemingly inhibitory components from DM52. Similarly, we observed an increased growth rate moving from DM16 to DM13, likely attributable to removing the most inhibitory component in the DM16, ammonium chloride. Notably, the media formulations lead to differing inter-run variability in OD600 measurements as well, with DM52 having the highest inter-run variability; this may be explained by the complexity of the media that could have resulted in different distribution of metabolic flux across pathways between runs. DM24 resulted in the lowest variability. The different formulations also have different average lag times, as identified from the fit function; DM52 and DM13 had the shortest (both ∼1.15 h) and DM16 with the longest (∼3.19 h). DM24 had an average lag time of ∼2.3 h. The maximum OD600 achieved is similar across DM52, DM24, and DM13 (OD600 ∼1.2) but lower in DM16 (OD600 ∼0.7). All growth curve features across the media formulations can be found in Supplemental File 6. We concluded that of these formulations, DM24 is the best general purpose medium due to supporting the highest maximum growth rate, lowest inter-run variability, short lag time, high max OD600, and the relative ease and low cost of its preparation. As such, we prioritized use of DM24 in downstream metabolite secretion experiments.

### *i*GR384 enables accurate simulation of *in vitro* LGG growth phenotypes

Since the DM series is fully chemically defined, these media and the single-component dropout phenotypes can be directly modeled using FBA^57^ simulations to predict biomass production rate, a proxy for growth rate, using the biomass objective function^58^. While neither the AGORA nor MERLIN LGG models support biomass flux in any formulation in our DM series, *i*GR384 supported biomass production in all formulations (Figure 3A). The precise simulated biomass flux matched *in vitro* growth rate results particularly well for growth in DM24, as the *in silico* biomass flux was 0.27/h while the average *in vitro* growth rate was 0.29/h. In DM52, *i*GR384 overpredicts the *in vitro* growth rate, but this is expected since several DM52 components seem to have negative regulatory effects on growth *in vitro*. Regulation of metabolism is not directly represented in *i*GR384 and is out of the scope of most prior GEM reconstruction efforts^57^. *i*GR384 somewhat underpredicts the *in vitro* growth rate in DM16 and DM13.

**Figure 3.**
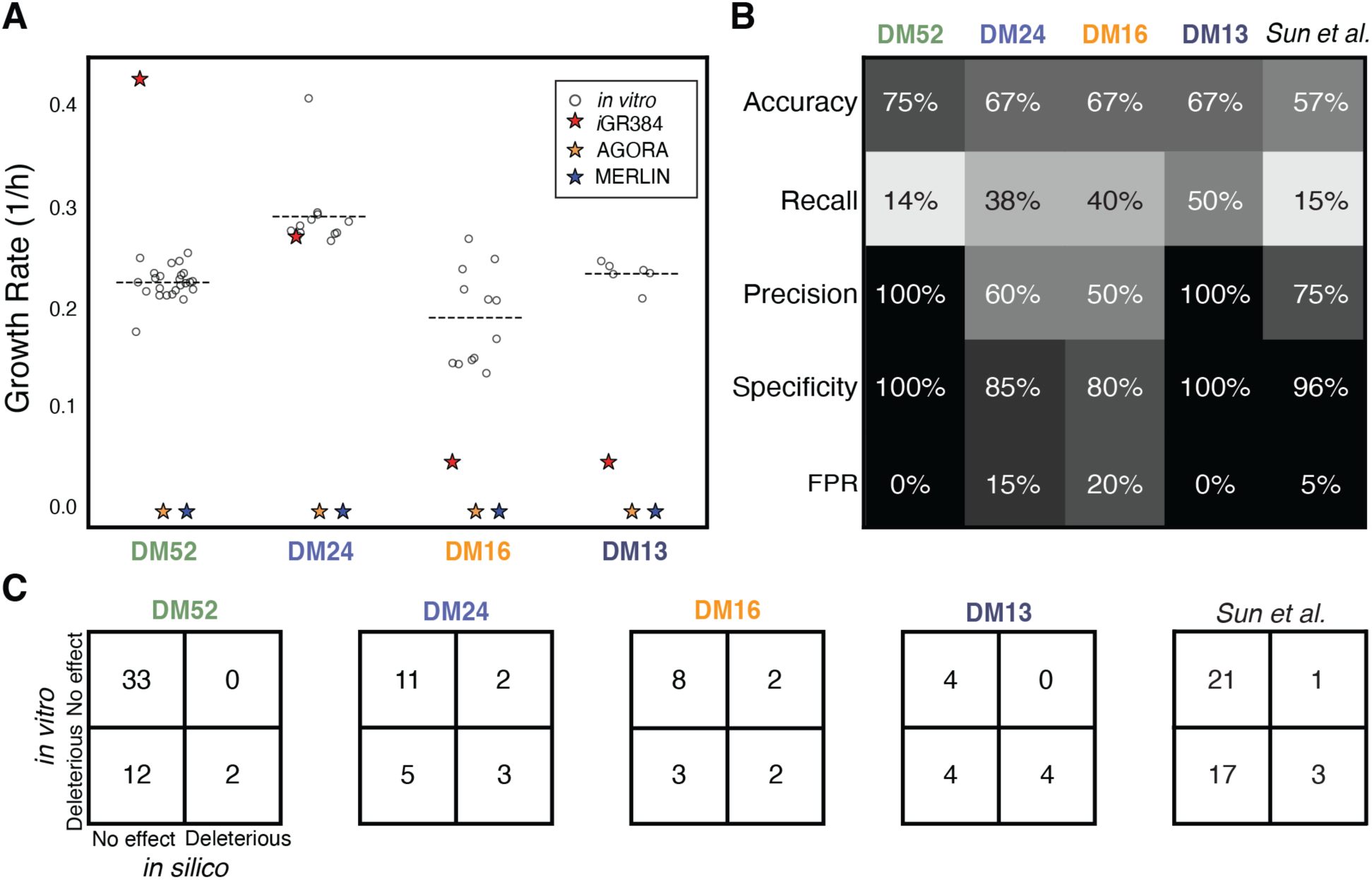
Validation of *i*GR384 against *in vitro* DM series and dropout growth data. *A. Comparison of simulated and measured LGG growth rates*. Circles indicate maximum specific growth rates from individual *in vitro* replicates, and dashed lines indicate the average across replicates. Stars indicate predicted maximum biomass flux from FBA simulations. *B. Performance of iGR384 in predicting dropout growth phenotypes.* Shading corresponds to 0 to 100% scale. FPR = false positive rate. *C. Confusion matrices of iGR384 in predicting dropout growth phenotypes.* Counts of (left to right, then top to bottom) true negatives, false positives, false negatives, and true positives are given with respect to dropout growth phenotypes from each background DM formulation.

In addition to comparing biomass flux and observed growth rate, traditional GEM validation also leverages gene essentiality studies^25,59–61^, which we replace with analogous single-component dropout experiments. In brief, nutrient dropouts are classified into “No Effect” or “Deleterious” categories from *in vitro* data. Then, these single-component dropouts are simulated and classified similarly based on biomass flux fold change relative to the corresponding full media formulation. Confusion matrix analysis follows (Figure 3C), and model performance metrics like accuracy, recall, precision, and specificity are calculated to assess how well the model recapitulates *in vitro* phenotypes (Figure 3B). *i*GR384 mostly recapitulates single component dropout phenotypes in all media formulations (Figure 3B), achieving at least 50% accuracy in all formulations. In DM52, *i*GR384 shows 75% accuracy, 100% precision and specificity, and 0% false positive rate (FPR). In DM24, *i*GR384 performs with 67% accuracy, 60% precision, 85% specificity, and 15% FPR. The model recall of deleterious phenotypes is low across all media, indicating that *i*GR384 often has alternative synthetic pathways to compensate for many nutrient losses. In addition, *i*GR384 performs much better than chance in the only other defined media for which single-component dropout growth data exist for LGG^35^. This formulation contains 44 components, including three not included in our DM series formulations, and differing concentrations of shared components. This last validation is especially notable as the previously published data was not considered during the reconstruction of *i*GR384 and therefore represents an especially stringent test.

### *i*GR384 enables analysis of LGG nutritional requirements

Defined minimal media are important tools in microbiology to understand the basic nutrient requirements for growth and as a nutrient environment baseline for perturbation experiments and metabolic engineering. Historically, minimal media have been determined through a laborious series of reductive defined media growth experiments, similar to our iterative single-component dropout experiments. GEMs offer a much faster and systematic approach to defining a minimal medium *in silico*. We used *i*GR384 to predict a defined simulated minimal medium for LGG (simMM). To achieve this, we determined the absolute required nutrients and any alternative nutrients for production of each individual biomass component and ATP maintenance (Figure 4A). These biomass component dependencies were used to guide the selection of compounds for a minimal medium. Glucose was not only a dependency for all biomass components and ATP maintenance but was one of only two dependencies for the exopolysaccharide, DNA, and RNA. Cysteine was a dependency for protein, lipid, and lipoteichoic acid. Glutamate and arginine are joint dependencies due to their participation in peptidoglycan, cofactor, and protein synthesis while not having sufficient *in vitro* data from the media dropouts to eliminate either one or the other. CoSO4 was included due to its essentiality in cofactor synthesis. Isoleucine was included due to its essentiality in protein synthesis. While aspartate was predicted as a dependency of all biomass components, it was ultimately excluded from the final minimal medium based on *in vitro* data showing that glutamate and aspartate are interchangeable and thus represents a known failure mode of *i*GR384 (Supplemental Figure 1). Together, each biomass component dependency is represented in our designed simulated minimal medium (simMM).

**Figure 4.**
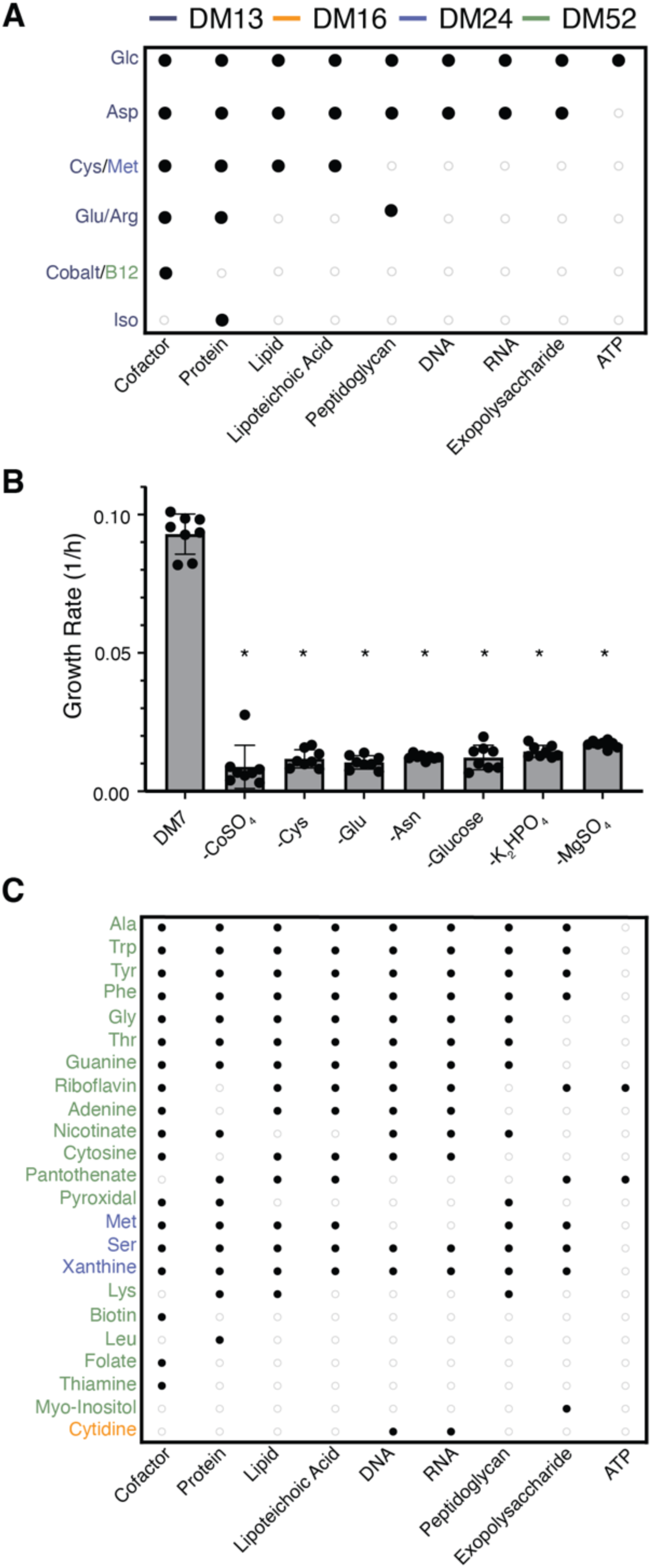
Model-guided determination of essential and supporting nutrients for biomass production. *A. Essential nutrients for biomass components.* Colors of nutrient names represent the most minimal medium condition in which they were predicted to be essential. These nutrients also remained essential in all more complex media formulations. *B. Model-informed minimal medium.* Average maximum specific growth rates from single-component dropout experiments (n=8) to prove essentiality of each component of minimal medium DM7. Significance of decreased growth rate relative to complete DM7 by unpaired t-test indicated by an asterisk (p-value < 0.001). *C. Biomass-supporting nutrients.* Colors of nutrient names represent the most minimal medium condition in which they were predicted as supporting the noted biomass components. These also remained supporting of noted biomass components in all more complex media formulations.

To achieve growth *in vitro*, we expanded upon simMM. Mineral salt requirements are notoriously hard to model in GEMs and were not directly represented in the biomass function, but our *in vitro* dropout data suggested that MgSO4 and K2HPO4 are essential salts for LGG growth and were retained in the minimal medium formulation. Additionally, asparagine is a protein dependency that also falls below the 20% growth rate reduction threshold when dropped-out of DM13 *in vitro* and was included in the model-informed minimal medium. Upon *in vitro* single-component dropout assessment of all nine of these compounds, arginine and isoleucine were determined to be non-essential (Supplemental Figure 2), leading to a model-informed true minimal medium called DM7, from which removal of any of the seven compounds leads to no growth of LGG (Figure 4B).

In addition to identifying required nutrients, we also identified nutrients that can support the production of biomass components (Figure 4C). We report 23 unique nutrients present in the DM series that can play a role in synthesis of essential biomass components even if not required. Together, these essential and supporting biomass dependencies define nutrient utilization capabilities of the LGG metabolic niche in the DM series.

### Theoretical characterization of indole-3-carboxaldehyde secretion requirements

Knowledge of LGG secretion of I3A comes from applied *in vivo* studies^12,14,62^ and is important for its therapeutic potential^4,13^. *Lactobacilli*, like *L. reuteri*, and other gut bacteria can catabolize tryptophan^63–65^, but the biochemistry of the final synthesis steps to I3A, including the enzymes responsible, remain unknown. To enable I3A secretion simulations using *i*GR384, we reconstructed a putative pathway for I3A synthesis in LGG based on indole pathways in related *Lactobacilli* (Supplemental Figure 3). Within this pathway, we identified all but two enzymes with orthologs in related organisms. We used *i*GR384 to predict I3A secretion as a function of biomass flux, generating production envelopes to characterize potential for I3A production in LGG using our DM formulations (Figure 5A-B). *i*GR384 predicts that I3A secretion flux is not growth coupled but can reach a theoretical maximum of ∼3.3 mmol/gDW/h in DM52 and ∼2.0 mmol/gDW/h in DM24. These maximum secretion rates are achieved only near 0 biomass flux, illustrating the direct tradeoff between biomass and I3A production. At observed *in vitro* growth rates, a theoretical maximum of ∼2.2 mmol/gDW/h and 0.0 mmol/gDW/h is predicted in DM52 and DM24, respectively. We then measured I3A production *in vitro* in DM52 via UPLC-MS after allowing LGG to ferment for 12, 24, and 48 hours and spanning both log and stationary growth phases (Supplemental Figure 4); however, no measurable I3A was detected. Since our *in vitro* experiments do not show detectable I3A secretion in DM52, there may be additional factors that contribute to I3A production *in vivo*, such as components of the much richer nutrient environment in the gut, quorum sensing or other regulatory effects.

**Figure 5.**
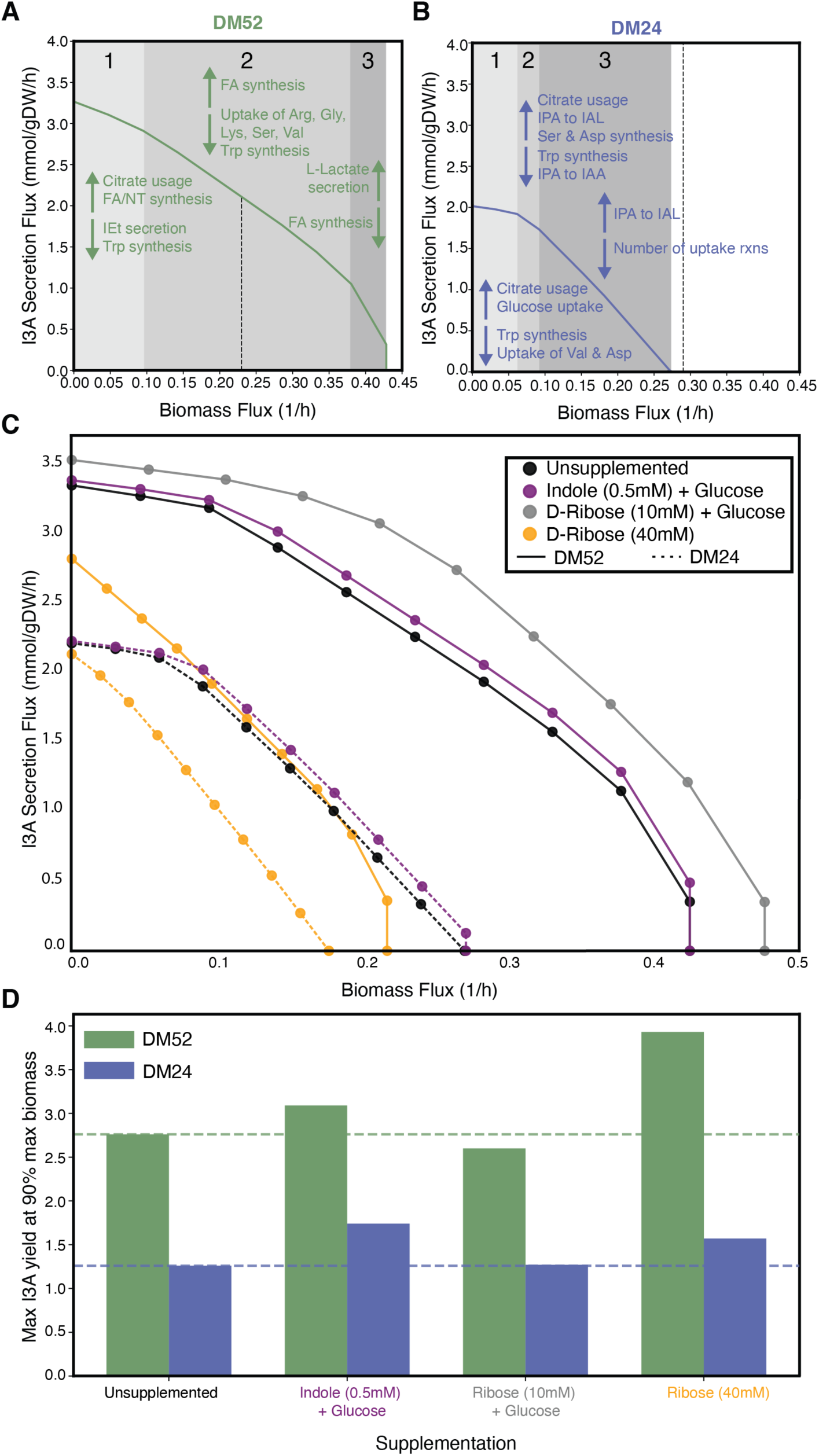
Predicted impact of media formulations on I3A secretion potential. *Simulated production envelopes for I3A secretion in (A) DM52 and (B) DM24.* Observed *in vitro* maximum specific growth rate indicated by dashed vertical line to represent corresponding expected potential I3A secretion. Grey highlighted areas denote distinct metabolic phases labeled with major shifts in metabolism that define them. IEt: indole-3-ethanol (tryptophol), IPA: indole-3-pyruvate, IAA: indole-3-acetate, and IAL: indole-3-acetaldehyde. All minimum I3A secretion fluxes were 0 for all biomass values. *C. Simulation of potential nutrient supplements to drive I3A secretion.* All minimum I3A secretion fluxes were 0 for all biomass values. Ribose supplementation in DM24 produced identical results to the unsupplemented control. *D. I3A yield from nutrient supplementation simulations.* Maximum I3A yield was calculated at 90% of the maximum possible biomass flux (I3A flux/biomass flux). Dashed lines represent the respective unsupplemented yield.

Although predicted secretion rates were low in both DM52 and DM24, metabolic phases of the production envelopes surrounding the observed *in vitro* growth rate may provide insight into the pathways and reactions that define the state of metabolism required for I3A secretion. These defining activities could be useful in future studies of I3A secretion. With relevance to I3A, tryptophan synthesis decreases around the *in vitro* growth rate in DM52 (Figure5A, Phase 2). In DM24, indole degradation products funnel more towards alternative product indole-3-acetaldehyde (IAL) rather than I3A around the *in vitro* growth rate (Figure5B, Phase 3). This provides insights into logical intervention points, like increasing tryptophan synthesis or driving indole degradation production towards I3A by decreasing flux through indole-3-pyruvate decarboxylase.

In addition to characterizing flux states with potential for I3A production in the base defined media, we applied *i*GR384 prospectively to explore the theoretical potential for nutrient supplementation to increase I3A secretion by LGG. Of the 90 metabolites with non-zero shadow prices^66–68^ for I3A secretion, three have evidence of crossing the bacterial cell membrane from supplementation experiments in other bacterial species and LGG genomic functional annotation of transporters represented in *i*GR384. These candidate I3A-promoting supplements include D-ribose (potential alternative carbon source) and indole (tryptophan precursor).

Indole increases maximum I3A yield marginally in both media conditions (Figure 5C-D)by increasing tryptophan degradation products upon satisfying tryptophan’s requirement in biomass. Ribose was tested both as an alternative and supplemental carbon source in addition to glucose. Ribose supplementation increases the maximum biomass flux and alters the shape of the production envelope in DM52, suggesting different pathway usage than the unsupplemented medium. However, ribose supplementation has no effect in DM24 (Figure 5C).

While it has been reported that LGG can grow on ribose^72^, the maximum biomass is significantly reduced in both media conditions. Full replacement of glucose with ribose in both DM52 and DM24 conditions alters the shape of the production envelopes and leads to increased I3A yield at 90% maximum biomass flux (Figure 5C-D), suggesting that metabolic states of LGG relying more heavily on ribose could increase I3A yield. This finding may have implications for dietary control of I3A secretion.

### Growth coupling of lactate secretion in LGG

Because lactate secretion by LABs is well understood, assessing this metabolic output serves as a benchmark for *i*GR384’s metabolic accuracy through comparative analysis between *in silico* and *in vitro* growth and lactate secretion experiments. We grew LGG in DM52 and DM24 in batch culture, paired with OD600 measurements via plate reader from the same inoculum with sampling every 2 h to guide duration of the growth curve (Supplemental Figure 5). Matching time point samples from the paired batch culture were collected for growth on solid medium to count colony-forming units (CFUs) and quantify lactate concentrations via HPLC (Figure 6A).

**Figure 6.**
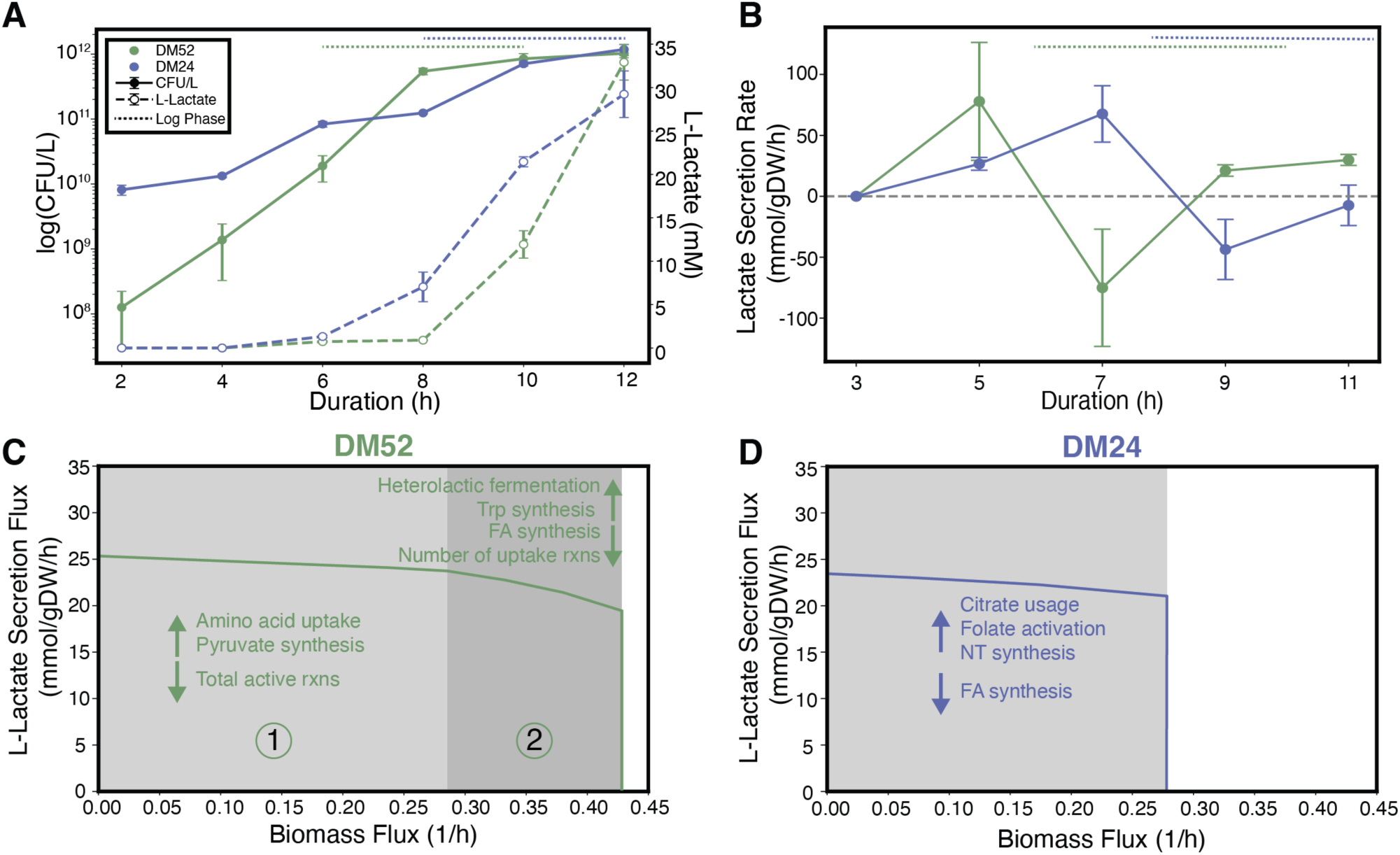
Differential lactate secretion in DM formulations. *A. in vitro growth and lactate secretion in DM formulations.* Average CFU/L measurements in logarithmic scale for three biological replicates reported on the left y-axis. Average lactate concentration across three biological replicates reported on the right y-axis. Error bars represent standard deviation. *B. Lactate secretion rates across growth curve.* Converted lactate secretion rates for all consecutive time points shown. Average and standard deviation plotted as the midpoint of consecutive time points used to calculate the rate. *Simulated production envelopes for lactate secretion in (C) DM52 and (D) DM24.* Grey-highlighted areas denote distinct metabolic phases in which characteristic pathway fluxes differ as indicated.

The difference in growth rate between DM52 and DM24 is mirrored in the lactate secretion measurements (Figure 6A) and supports the hypothesis that lactate secretion is growth-coupled in LGG in anaerobic conditions. It is well known that LABs can ferment sugars to lactic acid to generate ATP via substrate-level phosphorylation^73,74^. However, when assessing secretion rates across the growth curve, different fermentative strategies between the DM52 and DM24 are revealed (Figure 6B). In DM52, LGG has consistently positive lactate secretion rates in late log and stationary phase, indicating lactate fermentation occurring. In DM24, LGG ferments lactate in lag phase before switching to alternative pathways throughout log and stationary phase, evidenced by a positive to negative rate change. The complexity of DM52 may offer alternative strategies that contribute to growth during log phase and a return to canonical fermentation as those alternative strategies are exhausted. Overall, these findings suggest that LGG has different carbon resource allocation strategies in these different nutrient environments, as observed in other LABs^74,75^.

*i*GR384 predicts a maximum lactate secretion flux between ∼23 and ∼25 mmol/gDW/h in both DM52 and DM24 and a minimum of 0 mmol/gDW/h across the full range of biomass fluxes (Figure 6C-D). After converting *in vitro* secretion rates to comparable flux units (Supplemental File 7), we compared the measured lactate secretion rates in mid-log phase to the predicted secretion fluxes. In DM52, the lactate secretion rate in mid-log phase (at 9 h) was ∼20.0 mmol/gDW/h, close to *i*GR384’s theoretical maximum lactate secretion rate of 23 mmol/gDW/h. Systems-level analysis of reactions carrying flux in different metabolic phases on the production envelope (Figure 6C-D) gives insight into the different metabolic choices for carbon allocation in different DM formulations, allowing the model to suggest hypotheses that would be difficult to assess using *in vitro* approaches alone. For example, in both the DM52 and DM24, at observed growth rate, reactions involved in fatty acid synthesis have decreased flux. However, amino acid uptake supports the higher lactate secretion levels at lower biomass flux in the DM52, while usage of citrate increases as the biomass flux increases.

While the *in vitro* data is consistent with prior knowledge that lactate secretion is growth coupled in LABs, the LGG production envelopes indicate that lactate secretion is not fully growth coupled in *i*GR384. This is evidenced by the minimum lactate secretion rate being zero at all biomass flux values. Analyzing *i*GR384 to identify alternative pathways to lactate fermentation that can generate ATP revealed that amino acid and nucleotide metabolic pathways can also support ATP requirements for growth in these conditions (Supplemental Table 2). Fermentation-alternative ATP-generating pathways in LAB GEMs are not unique to *i*GR384. *i*NF517, a manually curated GEM for widely-studied lactate-producer *Lactococcus lactis* subsp. *cremoris* MG1363^76,77^ similarly exhibits high maximal lactate secretion in simulation conditions described by Flahaut *et al.* but can achieve maximum biomass flux with no lactate secretion as well, using similar amino acid metabolic pathways (Supplemental Figure 6). The network topology enabling non-fermentative ATP generation in these models may not be physiologically accurate under these growth conditions and might be resolved by contextualizing the models with condition-specific gene expression data^78^ or accounting, as knowledge permits, for other regulatory mechanisms limiting flux through non-fermentative pathways.

## Discussion

In this study, we developed critical resources for interdisciplinary study of *Lacticaseibacillus rhamnosus* GG (LGG) metabolism, including a validated genome-scale metabolic network reconstruction, a series of chemically-defined media of varying nutrient complexity including a minimal medium, and broad phenotypic growth data from single-component dropout experiments. We demonstrated how these resources can be integrated in the GEM reconstruction process and applied them to mechanistically study lactate fermentation and I3A secretion in LGG.

Commensal bacteria and probiotics have historically been studied primarily in an applied context, where they are introduced to a host system to observe potential prophylactic or therapeutic benefits. While these studies have led to some understanding of the connections between these microbes and effects on the host, relatively less is understood about the internal metabolism of these microbes themselves and the nutrient conditions required for them to exert these effects. The dearth of scientific literature on the individual metabolism of these taxa both highlights the need for mechanistic reconstructions such as GEMs to further study their metabolism and presents a challenge for traditional GEM reconstruction methods, which typically rely significantly on evidence from published literature. Even the most recent GEM of the most-well-studied model bacterium *Escherichia coli* MG1655, *i*ML1515^79^, has direct literature-based evidence for only ∼58% of its reaction content. This indicates the need for approaches and resources in the reconstruction of GEMs for such species.

Many Constraint-Based Reconstruction and Analysis (COBRA) studies approach this reconstruction problem by leveraging functional annotation of sequenced genomes to generate draft networks. While faster and requiring less extensive experimental data, resulting reconstructions are notoriously less reliable for detailed metabolic studies^32,33^. We illustrated that expanding upon the standard GEM reconstruction approach^25^ to include a consensus approach that leverages both toolkits to integrate experimental information can yield a useful reconstruction. We added merging steps after generating draft reconstructions by automated approaches, which decreases bias across representations of the same annotated genome. The merged draft reconstruction was manually curated based on experimental and literature evidence. Our reconstruction approach could be applied to many other organisms for which *in vitro* culture techniques and draft networks are available^26,30^. For example, *Fusobacterium nucleatum*, an important bacterium associated with colorectal cancer, has several automatically-generated draft reconstructions^26,82,83^, and lacks expansive mechanistic literature.

The integration of high-throughput defined media experiments into GEM reconstruction suggests a solution to overcome some of the resource limitations of understudied species and the painstaking manual steps required in the traditional reconstruction process. Biofoundaries coupled with artificial intelligence (AI) have been demonstrated to effectively enable autonomous microbiological laboratory experiments that explore defined media space to elucidate nutrient requirements analogous^80^ to those performed here^81^. Such AI-driven biofoundaries, if coupled with yet-to-be-developed next-generation GEM reconstruction algorithms, could perform the entire reconstruction process faster, with less manual input, and lead to higher-quality and more biologically accurate GEMs than those generated by current state-of-the-art automated reconstruction methods^26,30^ and perhaps improve GEMs even for well-studied species.

Development of the defined media series through single-component dropout experiments revealed insights into LGG’s nutritional requirements. We found that several amino acids, including cysteine, are essential to LGG growth. *Lactobacillus iners* is also a cysteine auxotroph, and it is well known that LABs commonly have amino acid autotrophies that contribute to their niche formation and community dynamics^84–87^. We also observed that LGG lacks a requirement for vitamins. Some *Lactobacilli* are vitamin auxotrophs, while others, like LGG shown here and other *Lacticaseibacillus* genus members, require cobalt for vitamin B12 synthesis but can otherwise synthesize all vitamins required for growth^88–91^.

Additionally, we evaluated essential and supporting roles of each component of the DM series for its contribution to biomass. These dependencies can be thought of as nutrient requirements defining a metabolic niche. Previous attempts have used GEMs to define metabolic niches based on the exchange fluxes needed to support biomass and secretion products^92^. While our current approach focuses on just the nutrient inputs defining a niche, we expand on this concept through comprehensive identification of not only essential but also supporting nutrients and resolve which specific biomass components depend on them. Coupled with secretion fluxes, this metabolic niche analysis approach could be extended to multi-species community modeling to study competitive and mutualistic relationships among species in a defined nutrient environment.

In addition to understanding niches, GEM-enabled nutrient environment optimization could be used as another lever to control target microbes’ participation in a microbial community and enhance integration of microbial GEMs into existing scaffolding for modeling whole humans^93^. Furthermore, the comprehensive identification of nutrient requirements–ranging from essential nutrients to compounds that improve specific biomass components, as characterized here for LGG using *i*GR384, could be coupled with potential for major secretion products to comprise a systematic definition for microbial metabolic niches. Establishing such metabolic niche definitions across a community, such as the gut microbiome, would provide a strong foundation for understanding and controlling community dynamics and host interactions.

Understanding factors affecting lactate production in LGG increases its potential in applications as a probiotic and food additive. We observe that dynamics of lactate fermentation vary in different media conditions. In the richest nutrient environment we tested LGG secretes less lactate overall than in a more minimal nutrient environment across all growth phases. In both media conditions, LGG also secretes more lactate during stationary phase than during mid-log growth phase, an encouraging property for probiotic applications. Understanding these dynamics is applicable to leveraging community interactions in the human gut microbiome for therapeutic purposes, since lactate can participate in metabolic cross-feeding interactions resulting in production of host-beneficial metabolites, such as butyrate^8,9,94^ and I3A^4,13,65^. Lactate also contributes to flavoring and vitamin stabilization^95,96^ in food products^97,98^.

In addition to evaluating lactate secretion, we also assessed LGG’s secretion of host beneficial I3A, concluding that LGG does not secrete detectable I3A levels in our DM series. Nevertheless, *i*GR384 serves as a powerful hypothesis generation tool; simulations revealed nutrient supplements that could be further tested. Indole has been explored in other species in the context of biofilm formation^69,70^. This suggests that I3A secretion may be tied to biofilm synthesis in LGG, a known participant in biofilms^99^. Another nutrient supplement, ribose, is predicted to increase biomass flux in LGG and may be a preferred carbon source in the context of I3A production. This could be explored to control I3A production by LGG *in vivo*.

The new tools and conceptual framework we introduce here for mechanistic understanding and control over LGG metabolism potentiate several future applications of *i*GR384, studying metabolic niches within microbiomes, and advancing methods for GEM reconstruction broadly. LGG is of significant therapeutic potential, and tools have been developed for its genetic manipulation^100–102^. GEMs have long been used to design bacterial strains^103–105^ with optimized secretion of target metabolites, which could be applied to LGG’s therapeutic secretions to effect microbial community dynamics and modulate gut-organ axis interactions.

## Materials and Methods

### Bacterial culture

For all growth experiments, *Lacticaseibacillus rhamnosus* GG (ATCC 53103) was grown at 37°C anaerobically (maximum 5-10ppm O2) in a Type C vinyl Coy chamber shaking at 150 rpm in various media conditions. LGG was recovered from 25% glycerol stocks stored at -80°C by fully thawing an entire 2 mL glycerol stock tube to inoculate in DM52 and culturing for 12 hours in 1:5 fresh DM52 media (total volume of 12 mL) to OD600 ∼1.5. Cultures were then diluted to OD600 of 0.05 to start experiments. For experiments using the DM13 and DM7 media, cells were washed three times by pelleting recovered culture at 10,000 *g* for 10 min, decanting, and replacing with 1X PBS before experimental inoculation. For all other experiments, experimental media were inoculated directly from recovery culture.

### Growth analysis

Growth curves were generated by both optical density measurement and colony counting approaches. A Cerillo Alto 96-well plate reader (ALT-1600) secured in the anaerobic chamber on a shaking incubator was used to measure OD600 of 200 μL of culture per well at 10-min intervals for 12 h. Growth curves were fit to OD600 data using ComBase’s DMfit^106^ tool, and max specific growth rate, R^2^ values, and other curve parameters were determined (Supplemental File 6). Curves were fit using the Baranyi-Roberts model^107^ where possible. Fitting algorithm and fit information for technical replicates are reported in Supplement File 6.

Growth curves accompanying HPLC results were determined using OD600. Additionally, colony-forming unit (CFU) counting was performed at specific timepoints, using a paired batch growth in parallel with the plate reader cultures described above. A 30 mL culture was used to aliquot technical replicates into the 96-well plate for OD600 measurements. 300 μL samples were collected from the 30 mL culture at 0 h, 2 h, 4 h, 6 h, 8 h, 10 h, and 12 h. At each time point, set dilutions were plated to ensure colony separation and sufficient colony counts for CFU/mL calculations. At 2 h, 1:100 and 1:1,000 dilutions were plated for both media formulations; 1:100 was counted for DM52, and 1:1,000 was counted for DM24. At 4 h, 1:1,000, 1:10,000, and 1:10,000 were plated; the 1:1,000 and 1:10,000 were counted for DM52 and DM24, respectively. At 6 h and 8 h, 1:10,000 and 1:100,000 were plated; 1:10,000 and 1:100,000 were counted for DM52 and DM24, respectively. At 10 h and 12 h, the 1: 100,000 and 1:1,000,000 dilutions were plated and the 1:1,000,000 plates counted for both the DM52 and DM24. All plates were inoculated with 100 μL diluted culture on Brain Heart Infusion broth (Remel, R452472) with 2% agar (Sigma-Aldrich 05039-500G). Plates were incubated aerobically at 37°C for 48 hours. Colonies were counted using Promega’s ColonyCount App and reviewed and corrected manually prior to calculating CFU/mL. The remaining 200 μL of collected sample was pelleted at 10,000 *g* for 10 min, and the supernatant was stored at -20°C for HPLC and UPLC measurements.

### Chemically-defined media and single-component dropout experiments

The initial set of compounds for defined media in this study contained 57 components, derived by merging the formulae of 3 different chemically defined media for *Lactobacilli* ^43–45^; the highest concentration was selected for redundant components. Five components were removed from the original 57-component list to generate our DM52 starting medium. These components were removed due to technical limitations with pH modulation (potassium phosphate monobasic), detergent activity (Tween 80) and solubility (2-deoxyuridine), being an experimental variable in the source paper (lipoic acid), and being an LGG secretion product of interest in this work (calcium lactate).

The media pH was measured and adjusted to 7.4 as needed with 1N NaOH prior to filter sterilization and overnight anaerobic degassing. Comprehensive single-component dropout experiments from DM52 were performed, replacing each component with the same volume of MilliQ water. Growth curves were derived using OD600 measurement and curve fitting as described above. Statistical significance of the difference in dropout max growth rates to that from full DM52 was assessed by unpaired t-test with p-value < 0.05. Dropout components with no significant decrease in growth rate were removed in formulating the subsequent DM. Some dropout components, such as aminobutyrate and valine, had results approaching significance but led to aberrant growth curve shapes that strongly influenced the computed max growth rate, leading to uncertainty about their effects. Such components were retained in the subsequent formulation. This procedure to reduce the media formulations was repeated iteratively, yielding DM24, DM16, and DM13 (Supplemental File 5).

### Measurement of secreted LGG metabolites

L-lactate was measured via high-performance liquid chromatography (HPLC) with UV-Vis in reserved supernatants from growth experiments, using a protocol adapted from a previous study measuring short-chain fatty acids^108^. Sodium L-lactate (98+%, Thermo Scientific, L14500.06) standards in DM52 were prepared by diluting 3:1 into methanol containing a final concentration of 1 mM internal standard 2-Ethylbutyric acid (98%, Alfa Aesar, A16989) prior to column filtration (Corning Costar Spin-X, 0.22). An Agilent HiPlex H column (300 x 7.7 mm, PL1170-6830) was used on an Agilent 1200 series HPLC. The column was run at 60°C with 5 mM H2SO4 with a flow rate of 0.6 mL/min for 48 min per sample injection. Compounds were detected at 210 nm. 20 μL of sample was injected, and autosampler temperature was maintained at 4°C. Each timepoint of the growth curve (0h, 2h, 4h, 6h, 8h, 10h, and 12h) consisted of 3 biological replicates, each injected once.

Indole compounds were detected by ultra-performance liquid chromatography (UPLC) using a protocol adapted from previous studies^4,109^. A 5 mM indole-3-carboxaldehyde (TCI, I0027), tryptophol (Sigma Aldrich, T90301-5G), and indole (99+%, Sigma Aldrich, I3408-25G) standard was prepared in DM52. A Waters XSelect Premier HSS T3 2.5 μm (2.1 x 150 mm, Part No. 186009856) column was used on a Waters Aquity UPLC system. The column was run at 40°C with a flow rate of 0.2 mL/min with a gradient of water + 0.1% formic acid (A) and methanol + 0.1% formic acid (B). The gradient was performed as follows (%A/%B): 0 min: 98/2, 2 min: 90/10, 6 min: 70/30, 10 min: 50/50, 14 min: 25/75, 15 min: 5/95, 17.5 min: 5:95, 17.51 min: 98:2, and 20 min: 98:2. Detection was performed using both the photodiode array (PDA) detector for UV-Vis at 260 nm and the Waters SQ Detector 2 with positive ion mode for mass spectrometry. Traces were identified using MS trace search using MassLynx version 4.2. Indole-3-carboxaldehyde was assessed at 145 g/mol, tryptophol at 162 g/mol, and tryptophan at 205 g/mol.

### Reconstruction of *i*GR384

#### Generation and merging of automated draft models

The AGORA LGG source reconstruction for genome assembly accession number ASM2650v1 was accessed via the AGORA 1.0 GitHub repository^110^. No alterations were made to this model prior to merging.

To generate the MERLIN LGG reconstruction, the genome sequence was retrieved from GenBank for assembly accession number ASM2650v1. Genome functional annotation was performed through Basic Local Alignment Search Tool (BLAST)^111^ searches against UniProtKB/TrEMBL and UniProtKB/Swiss-Prot^112^. Results from these alignments were used by MERLIN’s^113,114^ SamPler^115^ tool to assign annotations. The metabolic network was assembled automatically using MERLIN^113^. The metabolic data from the Kyoto Encyclopedia of Genes and Genomes (KEGG) database^36^ (including spontaneous, non-enzymatic reactions) was loaded and integrated with metabolic reactions from the LGG genome functional annotations. The genome sequence was analyzed using PSORTb 3.0^116^ to assign subcellular compartments to reactions. Transport reactions required for carrying metabolites through the compartments were identified using TranSyT^117^, available as a MERLIN plugin. Exchange reactions were automatically generated using MERLIN by integrating a set of reversible reactions for each metabolite found in the extracellular compartment. Gene-protein-reaction associations (GPRs) were formed automatically using MERLIN.

The biomass reaction for the MERLIN LGG reconstruction required manual curation, in addition to MERLIN tools. The “e-Biomass equation” tool^118^, implemented in MERLIN, was used to generate reactions for the synthesis of “e-Protein”, “e-DNA” and “e-RNA”. The GEMs of *Lactococcus lactis ssp. lactis* IL1403^119^, *Bacillus subtilis subsp. subtilis* 168^120^, *Lactobacillus plantarum*^121^, and *Lactobacillus casei*^122^ were used to inform “e-Protein”, “e-DNA” and “e-RNA”. Additional macromolecular metabolites, named as “e-Lipid”, “e-Exopolysaccharide”, “e-Peptidoglycan”, and “e-Lipoteichoic acid” were inferred from the mentioned reconstructions. Species-specific literature for *L. rhamnosus*^123–126^ was used to determine the exopolysaccharide, peptidoglycan, and average fatty acid compositions. An abstraction of the average fatty acid profile of LGG was represented as the synthesis of a pseudo-metabolite called “Fatty Acid”. Pseudo-metabolite “Fatty Acid” is a precursor to the synthesis of the macromolecule called “e-Lipid”. The lipoteichoic acid molecular structure was retrieved from published data from *L. rhamnosus*^127^. The lipid composition of *Lactobacilli* was retrieved from *Exterkate et al., 1971*^128^. The presence of cofactors and vitamins in the “e-Cofactors” was abstracted from reported evidence from literature^129,130^. To implement energy requirements, non-growth associated maintenance requirements for LGG were inferred from *L. lactis*^119,131^; then, the growth associated maintenance requirements were estimated through model fitting as described by Oliveira *et al.*^119^.

The entire set of reconstructed reactions was semi-automatically curated to ensure that the GEM was balanced and free of false network gaps. Reaction stoichiometry, mass balance, directionality and reversibility of all reactions as verified using MERLIN. Regarding reversibility, information was retrieved from the BiGG database^38^ and combined with a study by *Stelzer et al.*^132^ that analyzed reactions available in the KEGG database.

To merge draft reconstructions, all genes were converted to UniProtKB^133^ identifiers where possible. Any genes not existing in the UniProt database were retained as KEGG^36^ or MetaCyc^61^ identifiers. Reactions and metabolites were converted to their respective BiGG identifiers where possible, otherwise retained in KEGG or MetaCyc identifiers. Namespace mapping between reconstructions was performed first based on shared identifiers, then manually by reviewing descriptions and chemical formulae for reactions and metabolites. The merged reconstruction was seeded with matched reactions between AGORA and MERLIN reconstructions by merging those reactions to incorporate maximum information, and the unique reactions from each reconstruction were also added to the merged reconstruction. The curated biomass function from the MERLIN reconstruction was retained. This merged reconstruction was curated and refined to yield *i*GR384 (see below).

#### Iterative model evaluation and refinement using single-component dropout experimental phenotypes

Since all genes were initially identified and assigned to GPRs automatically, all genes in the model were manually reviewed for confident assignment to respective reactions based on functional annotation in their UniProtKB, KEGG, and MetaCyc entries.

Appropriate assignment was determined first by reaction equations present in database entries. Enzyme name and gene symbol were also considered and compared to information from literature for LGG or other closely related species. Inappropriate gene associations were removed from GPRs.

Gaps in the reconstruction after merging were identified using the “find_blocked_reactions” function in COBRApy^134^. Blocked reactions and pathways they participate in were reviewed for evidence in literature covering LGG and other related species and, where supported by literature evidence, additional pathway reconstruction was performed to fill the gaps in the model. Reactions not well-supported in literature but not refuted either were allowed to remain blocked in the model. Due to the literature-driven review of fatty acid metabolism in the generation of the MERLIN reconstruction and the modeling assumptions in the production of the preferred biomass function, gene-associated fatty acid synthesis reactions from the AGORA reconstruction were allowed to remain blocked, as they could represent a more detailed pathway to be fully functionalized in future versions of LGG models, but specific evidence did not exist to gap-fill beyond information already reflected in e_Fatty_Acid. All identified and remaining blocked reactions are noted in Supplemental File 2.

Many pathways required additional reconstruction to resolve gaps and add missing pathways to correct failure of the model to predict nutrient essentiality as identified by *in vitro* growth phenotypes in single-component dropouts from the DM52 medium. Due to sparse pathway-specific data in the LGG literature, a systematic approach was used to reconstruct putative pathways. KEGG pathways with LGG and related species’ gene annotation were used to identify the simplest putative path with the most coverage of reactions with gene associations to connect existing reconstruction reactions with strong supporting evidence, and reactions of the putative pathway were added to the reconstructions, including any non-gene associated intermediate reactions. Such related species that were prioritized, in order of priority, included: *Lacticaseibacillus casei* sp. and any other genus members*, Lactococcus lactis* sp.*, Bacillus subtilis* sp., and *Escherichia coli* sp. Candidate genes to support the added reactions for which an LGG gene was not associated in KEGG were identified by BLAST to search query genes associated with these reactions in related organisms against the LGG genome. Genes were accepted based on holistic hit review- including acceptable E-value, % identity, gaps, gene functional annotation consistency with query, and manual evaluation of alignment. Where no gene candidate was identified, the reaction was allowed to remain non-gene associated in the reconstruction.

After gap filling was completed, reaction directionality assessment, mass balancing, and loop reduction were performed according to standard reconstruction approaches^25^.

#### Expanding database cross-references, annotation, and supporting features

Additional unifying identifiers were extracted for all reactions, genes, and metabolites. Unifying databases included: KEGG^36^, MetaCyc^37^, MetaNetX^40^, ModelSEED^29^, RHEA^135^, E.C. Numbers^136^ for reactions, PubChem^137^ for metabolites, and RefSeq^138^ for genes. Subsystems for reactions were assigned based on other GEMs in the BiGG database, prioritizing for *i*JO1366^60^ (GEM for *Escherichia coli* K-12 MG1655) or *i*NF517^76^ (GEM for *Lactococcus lactis* subsp. *cremoris* MG1363) when multiple reconstructions contained the reaction. Systems Biology Ontology^139^ (SBO) terms for reactions were generated using SBOannotator^140^. Reaction evidence scores to inform confidence in their reconstruction were assigned based on a modified version of the standard reconstruction protocol^25^ (Supplemental Table 1). All annotations and supporting features are included in the tabular reconstruction file (Supplemental File 2).

#### Setting DM media conditions for *i*GR384 simulations

COBRApy version 0.29.1^134^ was used to perform all flux simulations. Defined media conditions in the DM series were set after converting concentrations of all components to exchange reaction bounds relative to the glucose concentration in the media (40 mM) and maximum glucose uptake rate (assumed to be 10 mmol/gDW/h). All exchange reaction bounds were confirmed to be above the solver (Gurobi^TM^) tolerance and were set using the “set_DM_usage.py” script provided on GitHub (Data Availability Statement).

#### Validation of model performance against *in vitro* experiments

Functionality of *i*GR384 in simulating growth phenotypes was validated using our newly reported *in vitro* single-component media dropout experiments. Maximum growth rates from dropouts were compared to that from growth in the respective full DM formulation. Individual components were classified as “No Effect” if the fold change was ≥0.8 and “Deleterious” if <0.8. The threshold of 0.8 was selected based on historical precedent to classify phenotypes in other media dropout and single-gene deletion studies^25,35,59–61,141,142^. To further support the propriety of this choice, we showed that a bimodal Gaussian mixture model (GMM) fitted to the fold changes across all *in vitro* dropout experiments resulted in a best-separating threshold of 0.94 (Supplemental Figure 7). This best-separating threshold was defined as the intersection of the two mixture-weighted Gaussian component densities, representing the point at which the posterior assignment probabilities for the two components are equal. Other thresholds can be derived from a GMM approach, including the unweighted component density. The intersection of the unweighted Gaussian component densities occurred at 0.73. Such intersection points have been used previously, especially within gene expression analysis^143^ and for non-biological thresholding problems with similar noisy data^144^. The chosen threshold of 0.8 lies within the range of 0.73 to 0.94 fold change.

For *in silico* simulations, full DM formulation conditions were initially set, and each dropout component was individually excluded by setting the uptake bound on the associated exchange reaction to 0. Flux balance analysis (FBA) was performed to determine maximum biomass flux in each dropout condition, and fold change versus the result in the full DM was calculated. Resulting fold changes were classified according to the same 0.8 threshold used to classify the *in vitro* results. The binarized *in vitro* and *in silico* dropout data was used to generate confusion matrices (Figure 3C) and model performance metrics calculated^25,145^.

#### Analysis of secretion product potential

Production envelopes^68^ were generated using the “PE_utils_usage.py” script provided on GitHub. In brief, FBA was performed to identify the maximum biomass rate, and values at regular intervals between 0 biomass and maximum biomass were selected. For each fixed biomass flux value, flux variability analysis (FVA^146^) was performed to determine the minimum and maximum achievable target metabolite secretion flux, represented by the appropriate exchange reaction. Raw *in vitro* data and calculations are presented in Supplemental File 7. *In vitro* L-lactate secretion rates, computed in mmol/CFU/L/h, were converted to comparable flux units using a conversion factor of 3.03E-13 gDW/CFU from LGG’s close relative *Lacticaseibacillus casei*^147^ taken as the average three measurements (Table 1).

**Table 1.** Summary of gDW conversion data from Nanjaiah et al, 2024 used to generate gDW/CFU conversion factor.

| gDW/L | CFU/mL |
| --- | --- |
| 3.95 | 10<sup>10.98</sup> |
| 3.36 | 10<sup>9.75</sup> |
| 4.6 | 10<sup>10.23</sup> |

#### Identification of alternative ATP-generating pathways in LAB models

Parsimonious FBA (pFBA)^148^ was performed to interrogate alternative pathways in *i*GR384 capable of generating ATP, aside from the expected lactate fermentation, after closing the lactate dehydrogenase reaction (i.e. setting upper and lower bounds on LDH_L to 0) in DM52 and DM24 conditions. This approach minimizes total network flux while optimizing the ATP maintenance objective function^148^, resulting in all non-zero fluxes contributing specifically to ATP generation. To analyze alternative ATP-generating pathways in *i*NF517, reaction bounds were set as previously reported^76^ with all exchanges given lower bounds of -10 mmol/gDW/h to represent a rich medium. Then, the same pFBA simulation was performed as described for *i*GR384.

#### Prediction of media supplements for increased I3A secretion

First, DM52 or DM24 conditions were set in *i*GR384, indole-3-carboxaldehyde (I3A) secretion was set as the objective function, and FBA performed. Then shadow prices were extracted from the solution object, and all intracellular components with non-zero shadow prices^66–68^ were curated for ability to cross the membrane. Components already present in a DM formulation were excluded as a supplementation candidate. Suitable supplementation concentrations for these compounds were determined from previous experiments in other bacterial species^69–71^, and uptake bounds were set relative to glucose concentration and assumed maximum glucose uptake rate as described above.

In *E. coli* MG1655, indole has been supplemented at many concentrations to explore its role in growth and biofilm formation^69,70^. From these studies, the median value of 0.5 mM was selected. Ribose can serve as an alternative carbon source to glucose for LGG, while less effective than for other *Lacticaseibacillus rhamnosus* strains^150^. We represented ribose supplementation using a nominal concentration of 10 mM ribose and also simulated completely replacing glucose with 40 mM ribose. Yield of I3A secretion per biomass flux unit was calculated at 90% of the maximum biomass flux for each simulated media condition.

#### Determining essential and supporting nutrients and developing a minimal medium

To determine the nutrients required for each biomass component, an artificial demand reaction was created for each biomass component pseudo metabolite in *i*GR384. With the DM13 media constraints set, each demand, along with the ATP maintenance function, was iteratively set as the objective function, and single component dropouts of each DM13 nutrient were assessed. Nutrients that brought objective flux below the solver tolerance were considered essential for the synthesis of that biomass component. For each subsequent higher-order media formulation (DM16, DM24, DM52), substitutions for the essential components were identified by first removing each essential component identified in a lower-order DM in turn and then removing each component of the current, higher-order medium to identify additional conditionally essential components leading to loss of objective flux. These conditionally essential components can act as substitutions for the lower-order essential component, restoring objective flux. The simulated minimal medium (simMM) was designed based on the essential components predicted in DM13.

To identify nutrients that were not absolutely required for, but could support production of, each biomass component including ATP maintenance, unique components from a higher-order DM were supplemented (with LB = -10) into the lower-order DM (e.g. DM16 component into DM13) and tested for increased biomass component flux. This process was performed for all subsequent DM pairs: DM13/DM16, DM16/DM24, and DM24/DM52.

Information regarding essential and supporting components were used to design a minimal medium following from observed LGG requirements from the DM13 *in vitro* dropout experiments. Finally, single-component dropout experiments as described above were performed to refine and confirm essentiality of the minimal medium components.

## Acknowledgements

We thank Dr. Peter Bellotti at the Albert Einstein College of Medicine Chemical Synthesis Core for assistance with UPLC and MS detection of indole compounds. We thank Dr. Sarah Wolfson for her assistance with HPLC approaches for lactate detection. We thank Carlos Madrid-Aliste for computational troubleshooting and local cluster management. We would like to thank Drs. Michelle Larsen and Christoph Kaleta for their support and guidance. GR would like to thank the members of the Chang and Kelly Labs for their feedback and useful discussions, particularly Renae Irving and Dr. Aleksandar Mihnev. GR was supported by the Albert Einstein College of Medicine Cellular, Molecular Biology, and Genetics T32 training grant and the PhRMA Foundation Predoctoral Fellowship in Drug Delivery. This research was supported in part by the National Institute of General Medical Sciences of the National Institutes of Health under award number R35GM151236 to RLC. This study was supported by the Portuguese Foundation for Science and Technology (FCT) under the scope of the strategic funding of UIDB/04469/2025 research unit, https://doi.org/10.54499/UID/04469/2025 and by LABBELS – Associate Laboratory in Biotechnology, Bioengineering and Microelectromechanical Systems, LA/P/0029/2020, https://doi.org/10.54499/LA/P/0029/2020.

## Author Contributions

**Gracelyn R. Richmond:** Conceptualization, Data Curation, Formal Analysis, Funding Acquisition, Investigation, Methodology, Project Administration, Resources, Software, Validation, Visualization, Writing-Original Draft, Writing- Review and Editing. **Emanuel Cunha:** Data Curation, Investigation, Methodology, Software, Writing-Original Draft, Writing-Review and Editing. **Libusha Kelly:** Methodology, Resources, Supervision, Writing- Review and Editing. **Óscar Dias:** Resources, Supervision, Writing- Review and Editing. **Roger L. Chang**: Conceptualization, Methodology, Funding Acquisition, Project Administration, Resources, Supervision, Validation, Visualization, Writing- Review and Editing

## Data Availability

All data, models, and code used in this study can be found on GitHub (https://github.com/RLChang-Lab/iGR384-LGG). The *i*GR384 model is also available on BioModels (<u>MODEL2603200002</u>).

